# Protein-protein interaction-Gaussian accelerated molecular dynamics (PPI-GaMD): Characterization of protein binding thermodynamics and kinetics

**DOI:** 10.1101/2021.09.27.461974

**Authors:** Jinan Wang, Yinglong Miao

## Abstract

Protein-protein interactions (PPIs) play key roles in many fundamental biological processes such as cellular signaling and immune responses. However, it has proven challenging to simulate repetitive protein association and dissociation in order to calculate binding free energies and kinetics of PPIs, due to long biological timescales and complex protein dynamics. To address this challenge, we have developed a new computational approach to all-atom simulations of PPIs based on a robust Gaussian accelerated molecular dynamics (GaMD) technique. The method, termed “PPI-GaMD”, selectively boosts interaction potential energy between protein partners to facilitate their slow dissociation. Meanwhile, another boost potential is applied to the remaining potential energy of the entire system to effectively model the protein’s flexibility and rebinding. PPI-GaMD has been demonstrated on a model system of the ribonuclease barnase interactions with its inhibitor barstar. Six independent 2 μs PPI-GaMD simulations have captured repetitive barstar dissociation and rebinding events, which enable calculations of the protein binding thermodynamics and kinetics simultaneously. The calculated binding free energies and kinetic rate constants agree well with the experimental data. Furthermore, PPI-GaMD simulations have provided mechanistic insights into barstar binding to barnase, which involve long-range electrostatic interactions and multiple binding pathways, being consistent with previous experimental and computational findings of this model system. In summary, PPI-GaMD provides a highly efficient and easy-to-use approach for binding free energy and kinetics calculations of PPIs.

## Introduction

Protein-protein interactions (PPIs) play key roles in many fundamental biological processes, including cellular signal transduction, immune responses and so on.^1^ Moreover, PPIs are implicated in the development of numerous human diseases and serve as important drug targets.^2^ Recent work has shown that developing PPI inhibitors represents a novel strategy for discovery of drugs with new mechanisms and a number of PPI inhibitors have been introduced to the market.^2-3^ Therefore, it is critical to investigate mechanisms of PPIs in both basic biology and applied medical research.

Experimental techniques^4^ including X-ray crystallography, nuclear magnetic resonance (NMR) and cryo-electron microscopy (cryo-EM) have been utilized to determine structures of protein-protein complexes. Recent years have seen dramatic increase in the number of experimentally determined protein-protein complex structures.^5^ However, it is still rather time-consuming and resource-demanding to obtain such experimental structures. Additionally, these experimental structures often capture only static pictures of protein-protein binding in the low-energy states. The very important intermediate conformational states with higher energy and dynamic mechanisms of the PPIs are still challenging to probe from the experiments.

A number of computational methods have been developed to explore PPIs and their binding mechanisms, including the widely used protein-protein docking^6^ and molecular dynamics (MD) simulations.^7^ The commonly used approaches for protein-protein docking include template-based docking such as HDOCK,^8^ GRAMM,^9^ GalaxyTBM^10^ and HADDOCK,^11^ and “ab initio” docking such as SwarmDock,^12^ ClusPro,^13^ GalaxyPPDock^10^ and MDockPP.^14^ Template-based docking remains the most accurate and widely used approach if a reliable template is available. Although many advanced ab initio approaches have been proposed, their accuracy could be still limited due to high degrees of freedom and protein flexibility during binding.^6^ While protein binding structures and free energies have been routinely predicted by docking, protein binding kinetics cannot be obtained from docking.

MD is a powerful technique for all-atom simulations of biomolecules to explore their structural dynamics, thermodynamics and kinetic properties.^7^ MD simulations have been applied to refine protein-protein docking poses.^15^ Additionally, MD has been widely applied to explore protein-protein binding mechanisms in atomistic detail.^16^ In this context, PPIs exhibit unique features, being distinct from the extensively studied protein-ligand interactions. The protein-protein binding affinity is often stronger than that of protein-ligand interactions. Protein-protein binding and unbinding processes often occurred in significantly longer time scales. Particularly, protein-protein dissociation process could take place over minutes, hours, and even days. Conventional MD (cMD) simulations have successfully captured fast binding of barstar to the barnase and revealed critical roles of water in the binding process.^17^ Despite remarkable advances, it remains challenging to sufficiently sample PPIs through cMD simulations, especially for the slow protein dissociation processes. Even with the specialized supercomputer ANTON, it is difficult to model protein-protein dissociation in hundreds-of-microsecond cMD simulations.^17b^

Enhanced sampling methods, including the Steered MD,^18^ Umbrella Sampling,^18-19^ Metadynamics,^16b, 20^ Weighted Ensemble,^21^ Replica Exchange MD,^15a^ Tempered Binding^17b^ and Markov State Models (MSMs),^22^ have been applied to improve sampling of PPIs. Such methodological advances have enabled simulations of millisecond or longer time scale processes, including protein-protein unbinding. For example, Umbrella Sampling was performed by Joshi et al. to delineate the barstar-barnase dissociation pathways and map the associated free energy landscape.^19^ Parallel tempering Metadynamics simulations with well-tempered ensemble (PTMetaD-WTE) and two carefully chosen collective variables captured both binding and unbinding processes of the insulin dimer.^8b, 13^ Weighted Ensemble with a total amount of ∼18 μs cMD simulations was implemented to investigate the mechanism of barstar binding to barnase.^21^ The predicted binding rate constant (k_on_) agreed well with the experiments.^21^ Pan et al.^17b^ developed the Tempered Binding method and captured repetitive association and dissociation events for five diverse protein–protein systems. Plattner et al.^22^ built MSM of barstar–barnase binding and unbinding processes with adaptive high-throughput cMD simulations. The obtained model revealed experimentally consistent intermediate structures, binding energetics and kinetics. However, such success was achieved by accumulating about two millisecond cMD simulations, requiring extraordinary computational resources. Therefore, while enhanced sampling methods have greatly expanded our capabilities in modeling PPIs, the above approaches remain computationally expensive for characterizing protein binding thermodynamics and kinetics. Compared with more extensively studied protein-ligand binding,^23^ enhanced sampling of PPIs is still largely underexplored.

Gaussian accelerated molecular dynamics (GaMD) is another enhanced sampling technique that works by adding a harmonic boost potential to smooth biomolecular potential energy surface.^24^ It greatly reduces system energy barriers and accelerates biomolecular simulations by orders of magnitude.^24b, 25^ Because the boost potential usually exhibits a Gaussian distribution, biomolecular free energy profiles can be properly recovered through cumulant expansion to the second order.^24b^ GaMD provides unconstrained enhanced sampling without the requirement of predefined reaction coordinates or collective variables. For PPIs, spontaneous binding of a G protein mimetic nanobody to a muscarinic G-protein-coupled receptor (GPCR) was successfully captured in previous GaMD simulations lasting for ∼4.5 μs.^26^ However, protein unbinding over longer time scales is still beyond the reach of normal GaMD simulations.^24a,26^ Notably, Paul et al. ^27^ introduced a new accelerated MD (aMD) approach, in which a boost potential was selectively applied on the van der Waals interactions between the protein MDM2 and its binding partner PMI. The new approach significantly increased chances to observe dissociation of the PMI from MDM2.

Building on GaMD, we have developed a new PPI-GaMD approach to explore the PPIs, in which the interaction energy potential of protein binding partners (both electrostatic and van der Waals interactions) is selectively boosted to facilitate protein dissociation. In addition, another boost is simultaneously applied on the remaining potential energy of the system to facilitate rebinding of the proteins. Moreover, we demonstrate PPI-GaMD on the dissociation and binding simulations of the barstar to barnase. The reason behind choosing this system is the availability of a wealth of experimental^3, 28^ and computational data^17, 19,21-22^ from extensive studies of this model system. To highlight efficiency of our PPI-GaMD approach, we focus on sampling the protein unbinding and rebinding events and predicting the protein binding free energy and kinetic rates. Remarkably, repetitive dissociation and association of barstar to barnase were observed in 2μs PPI-GaMD simulations. Protein binding free energies and kinetic rate constants calculated from PPI-GaMD simulations agreed excellently with the experimental data. Furthermore, PPI-GaMD simulations provided important insights into the binding mechanism of barstar to barnase at an atomistic level. We expect PPI-GaMD to be valuable for exploring rare events of protein dissociation and binding and calculating both the protein binding thermodynamics and kinetics.

## Methods

### Protein-Protein Interaction-Gaussian accelerated Molecular Dynamics (PPI-GaMD)

Based on GaMD^17b^, a new PPI-GaMD approach is developed here for further improved sampling of PPIs. We consider a system of ligand protein *L* binding to a target protein *P* in a biological environment *E*. The system comprises of *N* atoms with their coordinates 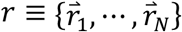 and momenta 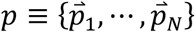. The system Hamiltonian can be expressed as:

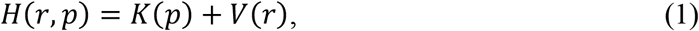

where *k*(*p*) and *V*(*r*) are the system kinetic and total potential energies, respectively. Next, we decompose the potential energy into the following terms:

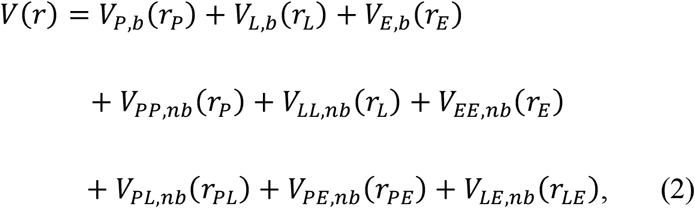

where *V*_*P,b*_, *V*_*L,b*_ and *V*_*E,b*_ are the bonded potential energies in protein *P*, protein *L* and environment *E*, respectively. *V*_*PP,nb*_, *V*_*LL, nb*_ and *V*_*EE, nb*_ are the self non-bonded potential energies in protein *P*, protein *L* and environment *E*, respectively. *V*_*PL, nb*_, *V*_*PE, nb*_ and *V*_*LE, nb*_ are the non-bonded interaction energies between *P-L, P-E* and *L-E*, respectively. According to classical molecular mechanics force fields^29^, the non-bonded potential energies are usually calculated as:

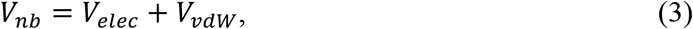

where *V*_*elec*_ and *V*_*vdw*_ are the system electrostatic and van der Waals potential energies. The interaction energy between the protein binding partners is *V*_*PL, nb*_ (*r*_*PL*_). In PPI-GaMD, we add boost potential selectively to the protein-protein interaction energy according to the GaMD algorithm:^17b^

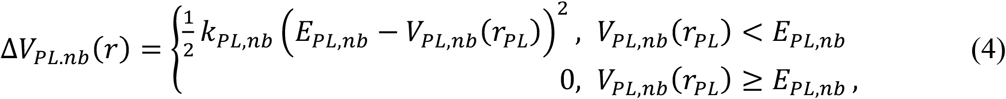

where *E*_*PL,nb*_ is the threshold energy for applying boost potential and *k*_*PL,nb*_ is the harmonic constant. The PPI-GaMD simulation parameters are derived similarly as in the previous GaMD algorithm.^17b^ When *E* is set to the lower bound as the system maximum potential energy (*E=V*_*max*_), the effective harmonic force constant *k*_0_ can be calculated as:

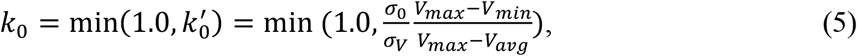

where *V*_*max*_, *V*_*min*_, *V*_*avg*_ and *σ*_*v*_ are the maximum, minimum, average and standard deviation of the boosted system potential energy, and *σ*_0_ is the user-specified upper limit of the standard deviation of Δ*V* for proper reweighting. The harmonic constant is calculated as 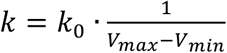 with 0 < *k*_0_ ≤ 1. Alternatively, when the threshold energy *E* is set to its upper bound 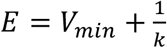, *k*_0_ is set to:

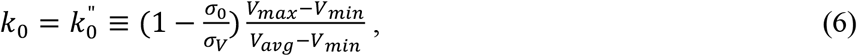

if 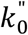 is found to be between *0* and *1*. Otherwise, *k*_0_ is calculated using Eqn. (5).

In addition to selectively boosting the interaction energy between proteins P and L, another boost potential is applied on the remaining potential energy of the system to enhance conformational sampling of the proteins and facilitate protein diffusion and rebinding. The second boost potential is calculated using the total system potential energy other than the interaction potential between the proteins as:

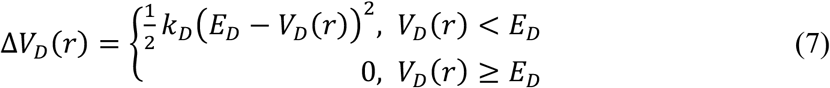

Where V_D_ is the total system potential energy other than the interaction potential between the proteins, E_D_ is the corresponding threshold energy for applying the second boost potential and k_D_ is the harmonic force constant. This leads to dual-boost PPI-GaMD with the total boost potential Δ*V*(*r*) = Δ*V*_*PL, nb*_ (*r*_*PL*_) + Δ*V*_*D*_(*r*). PPI-GaMD is currently implemented in the GPU version of AMBER 20,^30^ but should be transferable to other MD programs as well.

As noted above, the idea of selectively boosting the interaction energy between proteins is encouraged from previous aMD^27^ and Tempered Binding^17b^ studies. The aMD with selectively boosting the van der Waals interactions between the protein and peptide could significantly accelerate the slow peptide unbinding process.^27^ Besides, Tempered Binding^17b^ significantly accelerates the slow protein dissociation process by dynamically scaling electrostatic and van der Waals interactions between different groups of protein atoms by a factor, λ, which is updated among a ladder of discrete values, λ_i_. However, the current PPI-GaMD approach is different in terms of the following. First, the total non-bonded potential energy of both electrostatic and van der Waals interactions between proteins is boosted selectively. Second, another boost is applied to the remaining potential energy of the system to model system’s flexibility and facilitate the protein rebinding. Finally, the PPI-GaMD boost potentials are applied systematically according to the GaMD formula, which ensure proper reweighting of the simulations to recover the original protein binding thermodynamics and kinetics. It is worth noting that PPI-GaMD is also different from previous GaMD algorithms, including standard GaMD in which the boost potential was applied to the system dihedrals and/or total potential energy,^24b, 25a, 25b^ Ligand GaMD (LiGaMD) in which the non-bonded potential energy of the bound ligand is selectively boosted^31^ and peptide GaMD (Pep-GaMD) in which the essential potential energy of the highly flexible peptide is selectively boosted.^32^ In contrast, the total non-bonded interaction potential energy between the protein binding partners is selectively boosted in PPI-GaMD to enhance PPIs.

### Protein binding free energy calculations from 3D potential of mean force

We calculate protein binding free energy from 3D potential of mean force (PMF) of ligand protein displacements from the target protein as the following:^31-33^

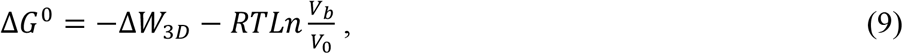

where *V*_0_ is the standard volume, *V*_*b*_ = ∫_*b*_ *e*^−*βw* (*r*)^ *dr* is the average sampled bound volume of the ligand protein with *β* = 1/*k*_*B*_*T, k*_*B*_ is the Boltzmann constant, *T* is the temperature, and Δ*W*_3*D*_ is the depth of the 3D PMF. Δ*W*_3*D*_ can be calculated by integrating Boltzmann distribution of the 3D PMF *W*(*r*) over all system coordinates except the x, y, z of the protein:

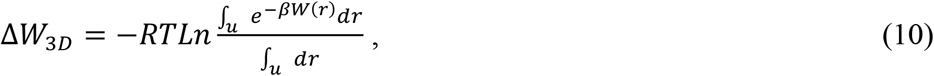

where *V*_*u*_ = ∫_*u*_ *dr* is the sampled unbound volume of the ligand protein. The exact definitions of the bound and unbound volumes *V*_*b*_ and *V*_*u*_ are not important as the exponential average cutoff contributions far away from the PMF minima.^33b^ A python script “PyReweighting-3D.py” in the *PyReweighting* tool kit (http://miao.compbio.ku.edu/PyReweighting/)^31, 34^ was applied for reweighting PPI-GaMD simulations to calculate the 3D reweighted PMF and associated protein binding free energies.

### Binding kinetics obtained from reweighting of PPI-GaMD Simulations

Reweighting of protein binding kinetics from PPI-GaMD simulations followed a similar protocol using Kramers’ rate theory that has been recently implemented in kinetics reweighting of the GaMD,^25c^ LiGaMD^31^ and Pep-GaMD simulations.^32^ Provided sufficient sampling of repetitive protein dissociation and binding in the simulations, we recorded the time periods and calculated their averages for the protein sampled in the bound (*τ*_*B*_) and unbound (*τ*_*U*_) states from the simulation trajectories. The *τ*_*B*_ corresponds to the protein residence time. Then, the protein dissociation and binding rate constants (*k*_off_ and *k*_on_) were calculated as:

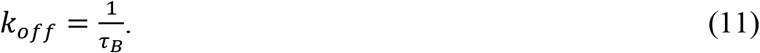

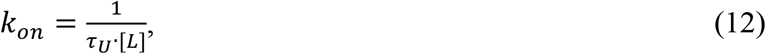

where [L] is the ligand protein concentration in the simulation system.

According to Kramers’ rate theory, the rate of a chemical reaction in the large viscosity limit is calculated as^25c^:

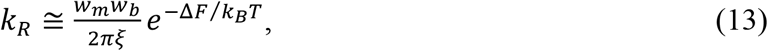

where *w*_*m*_ and *w*_*b*_ are frequencies of the approximated harmonic oscillators (also referred to as curvatures of free energy surface^35^) near the energy minimum and barrier, respectively, *ξ* is the frictional rate constant and Δ*F* is the free energy barrier of transition. The friction constant *ξ* is related to the diffusion coefficient *D* with *ξ* = *k*_*B*_*T*/*D*. The apparent diffusion coefficient *D* can be obtained by dividing the kinetic rate calculated directly using the transition time series collected directly from simulations by that using the probability density solution of the Smoluchowski equation.^36^ In order to reweight protein kinetics from the PPI-GaMD simulations using the Kramers’ rate theory, the free energy barriers of protein binding and dissociation are calculated from the original (reweighted, Δ***F***) and modified (no reweighting, Δ***F\****) PMF profiles, similarly for curvatures of the reweighed (*w*) and modified (*w*^∗^, no reweighting) PMF profiles near the protein bound (“B”) and unbound (“U”) low-energy wells and the energy barrier (“Br”), and the ratio of apparent diffusion coefficients from simulations without reweighting (modified, *D*^∗^) and with reweighting (*D*). The resulting numbers are then plugged into Eq. (13) to estimate accelerations of the protein binding and dissociation rates during PPI-GaMD simulations,^25c^ which allows us to recover the original kinetic rate constants.

### Simulation Protocols

PPI-GaMD simulations were demonstrated on the barnase-barstar system using the dual-boost scheme. The simulation system was constructed using crystal structure of the barnase-barstar complex (PDB ID: 1BRS)^37^ (chain B for the barnase and chain F superimposed to chain E for the barstar because some residues are missing in the chain E). Residues 40 and 82 in barstar were computationally restored to cysteine as in the wild-type protein. The missing hydrogen atoms were added using the *tleap* module in AMBER.^30^ The AMBER ff14SB force field^38^ was used for the protein. The system was neutralized by adding counter ions and immersed in a cubic TIP4P2015 water box,^39^ which was extended 18 Å from the protein-protein complex surface.

The simulation system was first energy minimized with 1.0 kcal/mol/Å^2^ constraints on the heavy atoms of the proteins, including the steepest descent minimization for 50,000 steps and conjugate gradient minimization for 50,000 steps. The system was then heated from 0 K to 310 K for 200 ps. It was further equilibrated using the NVT ensemble at 310K for 200 ps and the NPT ensemble at 310 K and 1 bar for 1 ns with 1 kcal/mol/Å^2^ constraints on the heavy atoms of the protein, followed by 2 ns short cMD without any constraint. The PPI-GaMD simulations proceeded with 14 ns short cMD to collect the potential statistics, 46 ns PPI-GaMD equilibration after adding the boost potential and then six independent 2000 ns production runs.

It provided more powerful sampling to set the threshold energy for applying the boost potential to the upper bound (i.e., *E* = *V*_min_+1/*k*) in our previous studies of ligand/peptide dissociation and binding using LiGaMD^31^ and Pep-GaMD.^32^ Therefore, the threshold energy for applying the boost potentials were all set to the upper bound in the PPI-GaMD simulations. In order to observe protein dissociation during PPI-GaMD equilibration while keeping the boost potential as low as possible for accurate energetic reweighting, the (σ_0P_, σ_0D_) parameters were set to (2.9 kcal/mol, 7.0 kcal/mol) in the final PPI-GaMD simulations. PPI-GaMD production simulation frames were saved every 0.2 ps for analysis.

The VMD^40^ and CPPTRAJ^41^ tools were used for simulation analysis. The 1D, 2D and 3D PMF profiles, as well as the protein binding free energy, were calculated through energetic reweighting of the PPI-GaMD simulations. The interface residues were defined as those from different protein subunits located within 10 Å between their Cα atoms in the X-ray complex structure.^17b^ Root-mean square deviation (RMSD) was calculated for the Cα atoms of the interface residues in the PPI-GaMD simulation frames relative to the X-ray structure after alignment. The protein interface distance was calculated between the center of masses (COMs) of the above defined interface residues from the different proteins. Protein interface RMSD and distance were chosen as reaction coordinates for calculating the PMF profiles. 2D PMF profiles of the interface RMSD and protein residue distances were calculated to analyze the protein binding pathways and important interactions. The bin size was set to 1.0 Å for these reaction coordinates. The cutoff for the number of simulation frames in one bin was set to 500 for reweighting of the 1D and 2D PMF profiles. The 3D PMF was calculated from each individual PPI-GaMD simulations of barstar binding to the barnase in terms of displacements of COM in the barstar interface residues from the COM of barnase interface residues in the X, Y and Z directions. The bin sizes were set to 1.0 Å in the X, Y and Z directions. The cutoff of simulation frames in one bin for 3D PMF reweighting was set to the minimum number below which the calculated global minimum of 3D PMF will be shifted. The protein binding free energies (*ΔG*) were calculated using the reweighted 3D PMF profiles.

## Results

### PPI-GaMD simulations significantly accelerated the barnase-barstar dissociation

All-atom PPI-GaMD simulations were performed on X-ray crystal structure of the barnase-barstar complex (**Fig. 1A**). Six independent 2,000 ns PPI-GaMD production simulations were obtained (**Table 1**). The PPI-GaMD simulations recorded average boost potentials of ∼35-36 kcal/mol with ∼6.3-6.4 kcal/mol standard deviation. The interface RMSD was calculated as a function of simulation time (**Fig. 1B**) to record the number of the barstar dissociation and association events (*N*_*D*_ and *N*_*B*_) in each of the 2,000 ns PPI-GaMD simulations. With close examination of the simulation trajectories, cutoffs of the interface RMSD for the unbound and bound states were set to >10 Å and <5.0 Å, respectively. Compared with the protein residence time determined from experiments (∼34.7 hours),^3^ PPI-GaMD significantly reduced the protein residence time to tens of nanoseconds by ∼12 orders of magnitude (**Fig. 1B**). Because of system fluctuations, we recorded only the binding and dissociation events lasting for more than 1.0 ns. Five binding and six dissociation events were observed in both Sim1 and Sim3. In Sim2, three binding and four dissociation events were captured. For the remaining simulations (Sim4 - Sim6), three binding and three dissociation events were recorded (**Table 1**). Therefore, repetitive barstar protein dissociation and rebinding were successfully captured in each of the 2,000 ns PPI-GaMD simulations (**Figs. 1B and S1**).

**Table 1.** Summary of PPI-GaMD simulations performed on binding of barstar to barnase. *ΔV* is the total boost potential. *N*_*D*_ and *N*_*B*_ are the number of observed barstar dissociation and binding events, respectively. *ΔG*_*sim*_ and *ΔG*_*exp*_ are the barnase-barstar binding free energies obtained from PPI-GaMD simulations and experiments, respectively.

| PDB | ID | Length (ns) | $\Delta V$ (kcal/mol) | $N_B$ | $N_D$ | $\Delta G_{sim}$ (kcal/mol) | $\Delta G_{exp}$ (kcal/mol) |
| --- | --- | --- | --- | --- | --- | --- | --- |
| 1BSR | Sim1 | 2000 | 35.74±6.35 | 5 | 6 | -17.79±1.11 | -18.90<sup>3</sup> |
|  | Sim2 | 2000 | 35.97±6.31 | 3 | 4 |  |  |
|  | Sim3 | 2000 | 35.78±6.36 | 5 | 6 |  |  |
|  | Sim4 | 2000 | 35.47±6.32 | 3 | 3 |  |  |
|  | Sim5 | 2000 | 36.06±6.40 | 3 | 3 |  |  |
|  | Sim6 | 2000 | 35.74±6.35 | 3 | 3 |  |  |

**Fig. 1.**
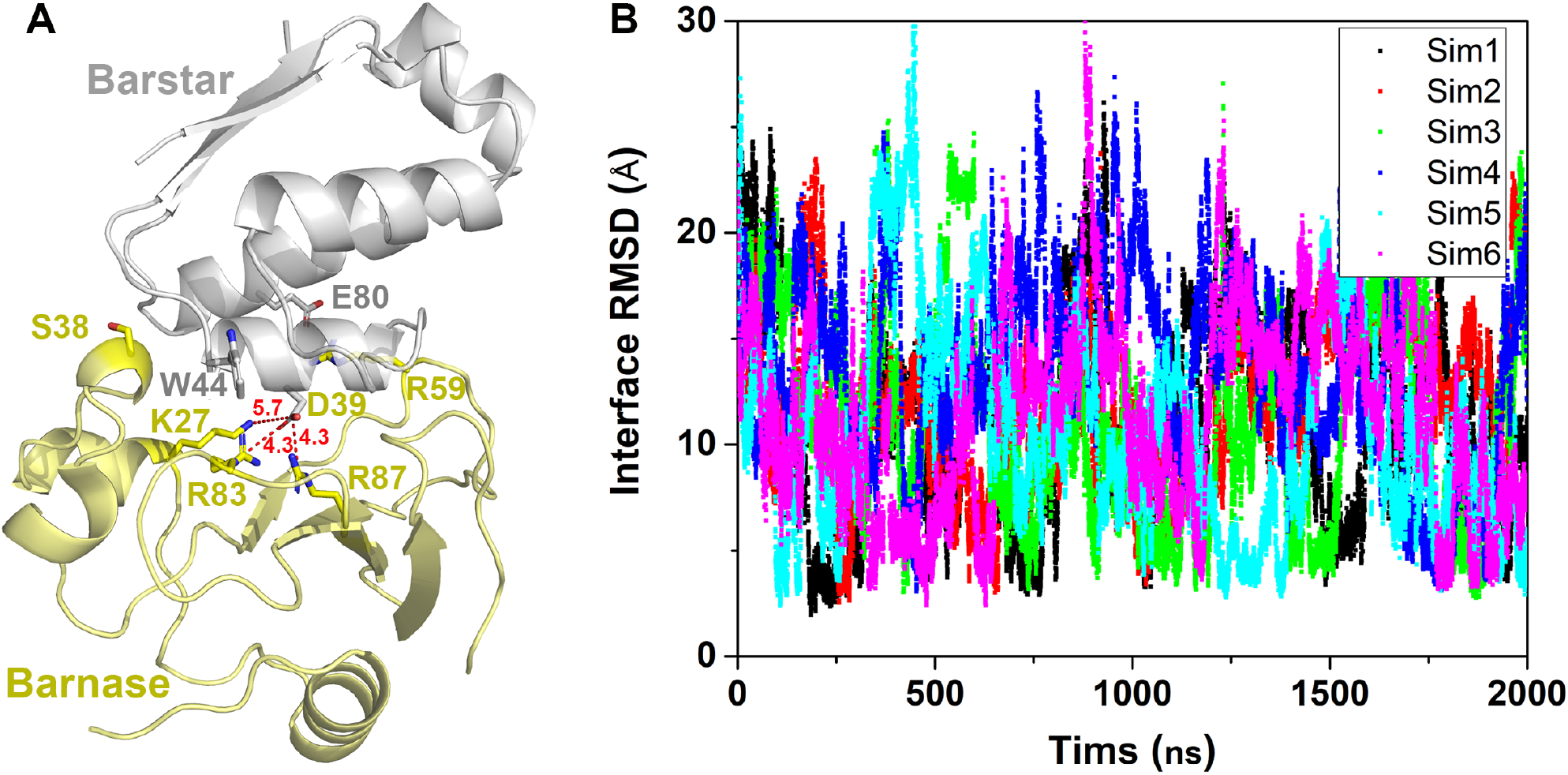
PPI-GaMD simulations captured repetitive dissociation and binding of barstar to barnase: (**A**) X-ray structure of the barnase-barstar complex. The barnase and barstar are shown in yellow and gray cartoons, respectively. Key barnase residues Lys27, Ser38, Arg59, Arg83 and Arg87 and barstar residues Asp39 and Glu80 are highlighted in sticks. Three salt bridges are shown in red dash lines with their corresponding distance values in angstrom; (**B**) Time courses of the protein interface root-mean-square deviation (interface RMSD) calculated from six independent 2,000 ns PPI-GaMD simulations. The interface residues are defined as those from different protein subunits located within 10 Å between their Cα atoms in the X-ray complex structure. Root-mean square deviation (RMSD) was calculated for the Cα atoms of the interface residues in the PPI-GaMD simulation frames relative to the X-ray structure after alignment.

### Multiple protein binding and dissociation pathways were identified from PPI-GaMD simulations

Free energy profiles were calculated from the PPI-GaMD simulations using the interface RMSD and the distance between barnase Arg59 and barstar Asp39 (denoted as BN:R59-BS:D39 distance) as reaction coordinates (**Fig. 2A**). One low-energy “bound” (B) conformational state was identified from the PPI-GaMD simulations, in which the interface RMSD and BN:R59-BS:D39 distance centered around (2.5 Å, 11.0 Å), being closely similar to the X-ray structure (**Fig. 2A**). Two low-energy “intermediate” states (I1 and I2) were also identified from the free energy landscape, in which the interface RMSD and BN:R59-BS:D39 distance centered around (7.5 Å, 6.0 Å) and (10 Å, 28 Å), respectively. Two low-energy “unbound” states (U1 and U2) were further identified, in which the interface RMSD and BN:R59-BS:D39 distance centered around (20.5 Å, 45.0 Å) and (27.5 Å, 20.0 Å), respectively.

**Fig. 2.**
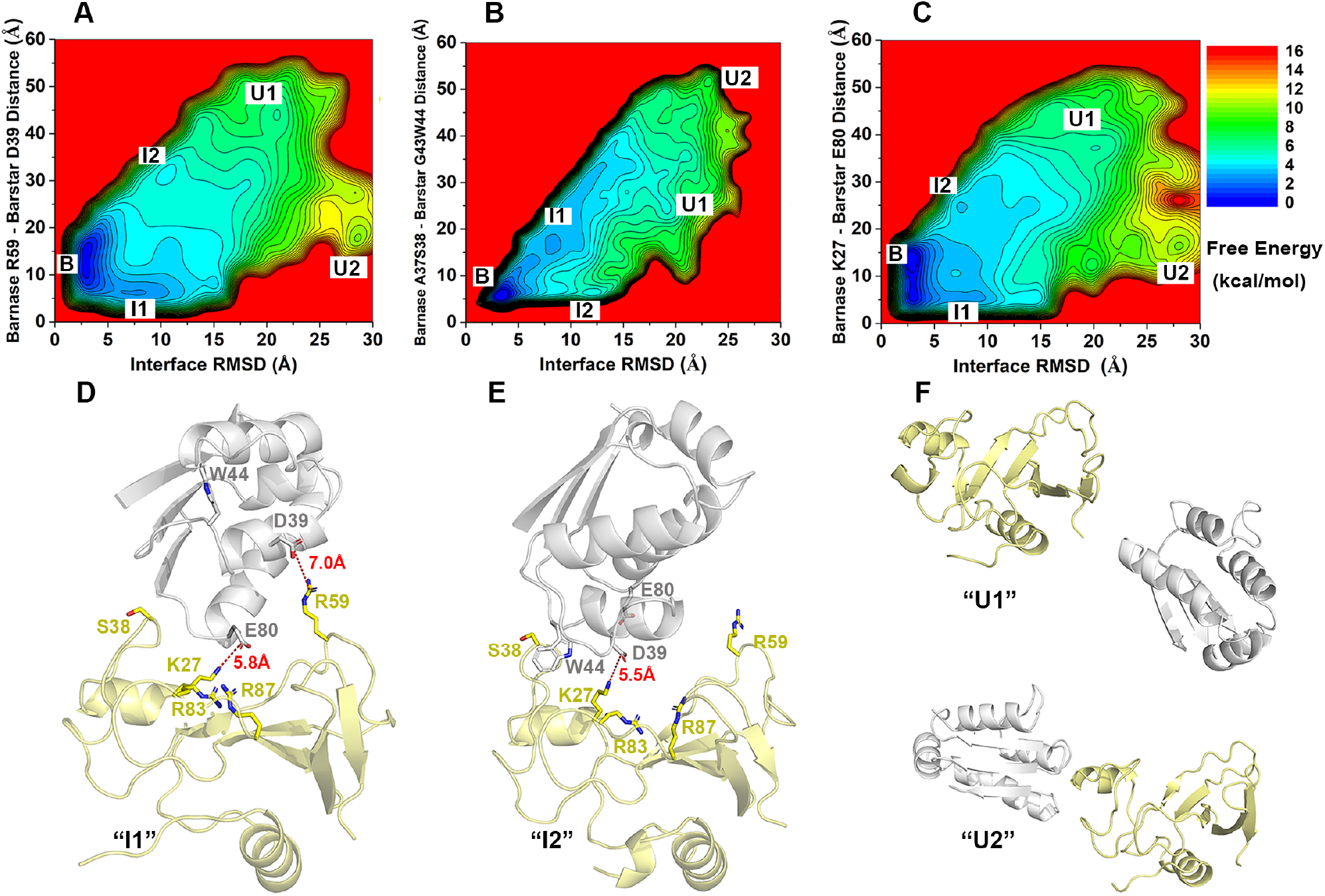
Free energy profiles and low-energy conformational states of barstar binding to barnase: (**A**) 2D PMF profiles regarding the interface RMSD and the distance between barnase Arg59 and barstar Asp39. (**B**) 2D PMF profiles regarding the interface RMSD and the distance between the center of masses (COMs) of barnase residues Ala37-Ser38 and barstar residues Gly43-Trp44. (**C**) 2D PMF profiles regarding the interface RMSD and the distance between barnase Lys27 and barstar Glu80. (**D**–**G**) Low-energy conformations as identified from the 2D PMF profiles of the (**D**) intermediate “I1”, (**E**) intermediate “I2”, (**F**) unbound “U1”, and (**F**) Unbound “U2”. Key barnase residues Lys27, Ser38, Arg59, Arg83 and Arg87 and barstar residues Asp39 and Glu80 are highlighted in sticks. Strong electrostatic interactions are shown in red dash lines with their corresponding distance values labeled in the intermediate “I1” (**D**) and “I2” (**E**).

The intermediate (I1 and I2) and unbound (U1 and U2) conformations were shown in **Figs. 2D-2G**. In the I1 state, the barnase Arg59 formed a salt bridge with the barstar Asp39 (**Fig. 2D**). Remarkably, positively charged residues (Lys27, Arg83 and Arg87) in barnase also formed electrostatic interactions with the negatively charged residue Glu80 in barstar (**Fig. 2D**). In the I2 state, the same positively charge residues in barnase (Lys27, Arg83 and Arg87) formed electrostatic interactions with the negatively charged residue Asp39 in barstar (**Fig. 2E**). Additionally, barnase Ser39 and barstar Trp44 were observed close distance in the I2 state (**Fig. 2E**). Therefore, the COM distance between the barnase Ala37-Ser38 and barstar Gly43-Trp44 (denoted as BN:A37S38-BS:G43W44 distance) and the distance between barnase Lys27 and barstar Glu80 (denoted as BN:K27-BS:E80 distance) were selected as additional reaction coordinates to calculate the free energy profiles (**Figs. 2B & 2C**). Similar “bound” (B), “intermediate” (I1 and I2) and “unbound” states (U1 and U2) were identified from the free energy profiles (**Figs. 2B and 2C**). In the intermediate I1 state, the BN:K27-BS:E80 distance and BN:A37S38-BS:G43W44 distance centered around 5.8 Å and 18.2 Å, respectively. In the intermediate I2 state, the BN:K27-BS:E80 distance and BN:A37S38-BS:G43W44 distance centered around 23.2 Å and 5.0 Å, respectively. With significant differences between the two intermediate states (**Figs. 2D& 2E**), barstar appeared to adopt two pathways for binding to the barnase. One was through electrostatic interactions between positively charged residues in barnase (Arg59, Lys27, Arg83 and Arg87) and negatively charged residues in barstar (Asp39 and Glu80) (**Fig. 2D**). The other was through hydrophobic interactions between BN:A37S38 and BS:G43W44 and electrostatic interactions between positively charged residues in barnase (Lys27, Arg83 and Arg87) and a negatively charged residue in barstar (Asp39) (**Fig. 2E**).

Structural clustering of barstar was applied to six individual PPI-GaMD trajectories (Sim1 to Sim6) to identify the representative binding pathways of barstar to barnase (**Fig. 3**). The structural clusters were reweighted to obtain their original free energy values ranging from 0.0 kcal/mol to 10.0 kcal*/*mol. The top 100 reweighted clusters were selected to represent the barstar binding pathways (**Fig. 3)**. Overall, binding of barstar to barnase occurred via two main binding sites, involving barnase residues Arg59 and Ala37-Ser38, respectively. Both sites contributed important interactions with barstar during its binding to barnase target site in the Sim1, Sim3, Sim4 and Sim5 (**Figs. 3A, 3C-3E**). While barstar bound to barnase mainly via interactions with barnase Arg59 in Sim2 (**Fig. 3B**). Barstar bound to barnase mainly via interactions with barnase residues Ala37 and Ser38 in Sim6 (**Fig. 3F**). These findings again revealed pathways for barstar binding to barnase. The two intermediate states were similar to those in the MSM obtained from ∼2 millisecond simulations.^22^ The two binding pathways were also similarly identified from Umbrella Sampling simulations.^19b^

**Fig. 3.**
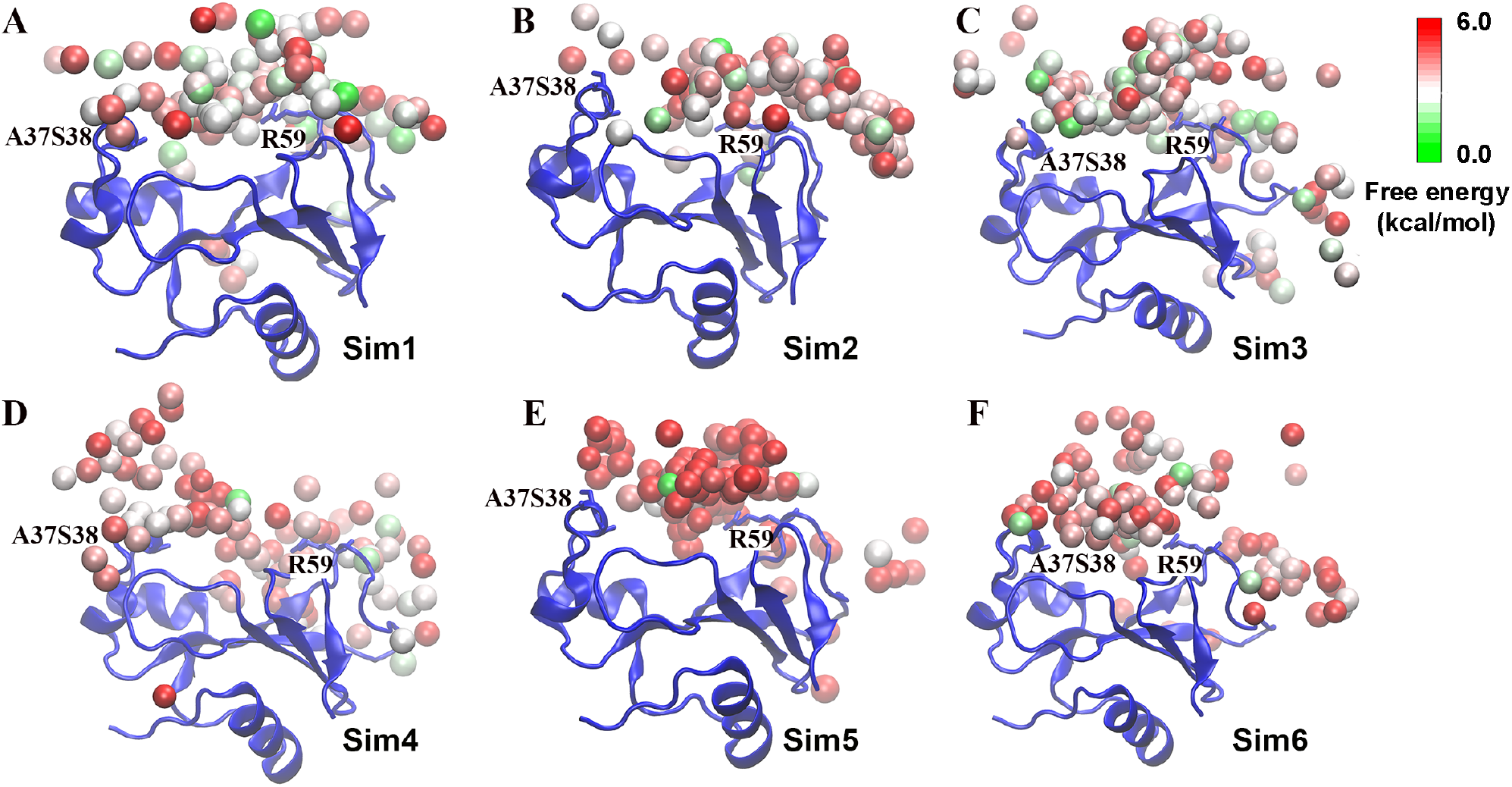
Binding pathways of barstar to barnase revealed from the (**A**) “Sim1”, (**B**) “Sim2”, (**C**) “Sim3”, (**D**) “Sim4”, (**E**) “Sim5” and (**F**) “Sim6” PPI-GaMD trajectories. The barstar interacted with barnase residues Ala37Ser37 or Arg59 in the intermediate conformations. Barnase is shown in blue ribbons. The barstar structural clusters (sphere) are colored by the reweighted PMF free energy values in a green (0.0 kcal*/*mol)-white-red (6.0 kcal*/*mol) color scale.

### Electrostatic interactions played an important role in the barstar binding/unbinding

Next, we examined key residue interactions during barstar binding to barnase (**Fig. 1**). In the X-ray structure (**Fig. 1A**), three salt bridges were found between barnase and barstar, i.e., BN:K27-BS:D39, BN:R83-BS:D39 and BN:R87-BS:D39. Therefore, the distances between these residues and the interface RMSD were further chosen as reaction coordinates to calculate 2D PMF profiles (**Fig. 4**). Five low-energy states were identified from the calculated 2D PMF profiles, including “bound” (B), “intermediate” (I1 and I2) and “unbound” (U1 and U2) (**Figs. 4A-4C**). In the “Bound” state, the (interface RMSD, BN:K27-BS:D39 distance), (interface RMSD, BN:R83-BS:D39 distance) and (interface RMSD, BN:R83-BS:D39 distance) centered around (2.5 Å, 7.5 Å), (2.8 Å, 8.2 Å) and (2.7 Å, 8.5 Å), respectively (**Fig. 4**). In the “I1” state, the (interface RMSD, BN:K27-BS:D39 distance), (interface RMSD, BN:R83-BS:D39 distance) and (interface RMSD, BN:R83-BS:D39 distance) centered around (7.5 Å, 15.0 Å), (7.8 Å, 17.2 Å) and (7.7 Å, 16.0 Å), respectively (**Fig. 4**). In the “I2” state, the (interface RMSD, BN:K27-BS:D39 distance), (interface RMSD, BN:R83-BS:D39 distance) and (interface RMSD, BN:R83-BS:D39 distance) centered around (11.0 Å, 5.8 Å), (10.5 Å, 9.0 Å) and (11.0 Å, 9.5 Å), respectively (**Fig. 4**). The representative intermediate conformational states (I1 and I2) were shown in **Fig. 2D & 2E**. In the “I1” state, barstar negatively charged residue Asp39 formed electrostatic interaction with positively charged residue Arg59 in barnase with BN:R59-BS:D39 distance at ∼7.0 Å (**Fig. 2D**). Additionally, barstar negatively charged residue Glu80 formed electrostatic interaction with positively charged residues Lys27, Arg83 and Arg87 in barnase, especially between BN:K27 and BS:E80 with a short distance of ∼5.8 Å (**Fig. 2D**). In the “I2” state, barstar negatively charged residue Asp39 formed electrostatic interaction with positively charged residues Lys27, Arg83 and Arg87, especially between BN:K27 and BS:D39 with a distance of ∼5.5 Å (**Fig. 2E**). Therefore, the above favorable electrostatic interactions in the “intermediate” states played an important role for the binding and unbinding of barstar, being highly consistent with the mutation experimental data.^28a, 42^ In summary, long-range electrostatic interactions played a key role in the binding/unbinding of barstar-barnase. The electrostatic interactions involving residues Lys27, Arg83 and Arg87 in barnase appeared to be an anchor to pull the barstar to its target binding site. Similar findings were obtained from previous MD simulations.^19b^

**Fig. 4.**
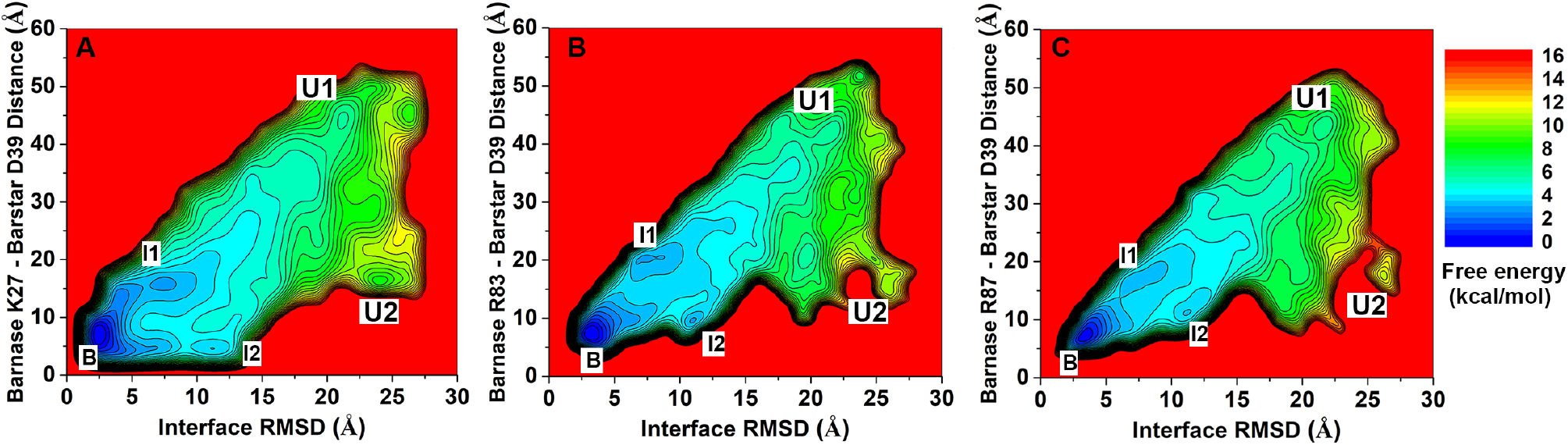
(**A**) 2D PMF profiles regarding the interface RMSD and the distance between barnase Lys27 and barstar Asp39. (**B**) 2D PMF profiles regarding the interface RMSD and the distance between barnase Arg83 and barstar Asp39. (**C**) 2D PMF profiles regarding the interface RMSD and the distance between barnase Arg27 and barstar Asp39.

### Protein binding free energy calculated from PPI-GaMD simulations agreed with experimental data

We calculated the binding free energy of barstar to barnase from each individual PPI-GaMD simulations based on 3D PMF of the displacement of barstar interface residues from the barnase interface residues in the X, Y and Z directions. The 3D PMF was energetically reweighted through cumulant expansion to the second order. We then calculated the protein binding free energy using the 3D reweighted PMF profiles. The average barstar binding free energy was −17.79 kcal/mol with a standard deviation of 1.11 kcal/mol, being highly consistent with the experimental value of −18.90 kcal/mol (**Table 1**).^3^ In summary, the binding free energy of barstar-barnase calculated from PPI-GaMD simulations agreed well with the experimental data. The prediction error in the PPI-GaMD simulations was 1.11±1.11 kcal/mol. Therefore, both efficient enhanced sampling and accurate protein binding free energy calculation were achieved through the PPI-GaMD simulations.

### Kinetics of barnase-barstar binding

With accurate prediction of the protein binding free energy, we further analyzed the PPI-GaMD simulations to determine the kinetic rate constants of barstar binding to barnase. We recorded the time periods for the barstar found in the bound (*τ*_B_) and unbound (*τ*_U_) states throughout the PPI-GaMD simulations. The barstar concentration was 0.0055 M in the PPI-GaMD simulation system. Without reweighting of the PPI-GaMD simulations, the barstar binding (*k*_*on*_*\**) and dissociation (*k*_*off*_*\**) rate constants were calculated directly as 8.02 ± 0.22 × 10^8^ M^-1^×s^-1^ and 1.44 ± 0.34 × 10^7^ s^-1^, respectively (**Table 2**).

**Table 2.** Comparison of kinetic rates obtained from experimental data and PPI-GaMD simulations for barstar binding to barnase. *k*_*on*_ and *k*_*off*_ are the kinetic dissociation and binding rate constants, respectively, from experimental data or PPI-GaMD simulations with reweighting using Kramers’ rate theory. *k*_*on*_*\** and *k*_*off*_*\** are the accelerated kinetic dissociation and binding rate constants calculated directly from PPI-GaMD simulations without reweighting.

| | Method | $k_{on}$<br>($M^{-1} \cdot s^{-1}$) | $k_{off}$<br>($s^{-1}$) | $k_{on}^*$<br>($M^{-1} \cdot s^{-1}$) | $k_{off}^*$<br>($s^{-1}$) |
| --- | --- | --- | --- | --- | --- |
| 1BRS | Experiment<sup>3</sup> | $6.0 \times 10^8$ | $8.0 \times 10^{-6}$ | - | - |
| | PPI-GaMD | $21.7 \pm 13.8 \times 10^8$ | $7.32 \pm 4.95 \times 10^{-6}$ | $8.02 \pm 0.22 \times 10^8$ | $1.44 \pm 0.34 \times 10^7$ |

Next, we reweighted the barstar-barnase PPI-GaMD simulations to calculate acceleration factors of the barstar binding and dissociation processes (**Table S1 & Fig. S1**) and recover the original kinetic rate constants using the Kramers’ rate theory (**Table 2**). The dissociation free energy barrier (Δ*F*_*off*_) decreased significantly from 6.91, 7.70, 7.60, 5.85, 9.40 and 7.72 kcal/mol in the reweighted PMF profiles to 1.41, 1.89, 1.91, 0.18, 1.67 and 1.68 kcal/mol in the modified PMF profiles for Sim1 to Sim6, respectively (**Table S1**). On the other hand, the free energy barrier for the barstar binding (Δ*F*_*on*_) decreased slightly from 1.06, 2.01, 4.80, 0.95, 1.52, and 1.16 kcal/mol in the reweighted profiles to 0.23, 0.10, 0.76, 0.16, 0.54, and 0.21 kcal/mol in the modified PMF profiles for Sim1 to Sim6, respectively (**Table S1**). The curvatures of the reweighed (*w*) and modified (*w*^∗^, no reweighting) free energy profiles were calculated near the protein bound (“B”) and unbound (“U”) low-energy wells and the energy barrier (“Br”), as well as the ratio of apparent diffusion coefficients calculated from the PPI-GaMD simulations with reweighting (*D*) and without reweighting (modified, *D*^∗^) (**Table S1**). According to the Kramers’ rate theory, the barstar association was accelerated by 27.67, 0.67, 0.43, 0.17, 0.12 and 0.15 times in Sim1 to Sim6, respectively. Remarkably, the barstar dissociation was significantly accelerated by 1.48 × 10^12^, 3.04 × 10^12^, 6.51 × 10^11^, 2.54 × 10^12^, 9.43 × 10^13^ and 1.26 × 10^13^ times in Sim1 to Sim6, respectively. Therefore, the average reweighted *k*_*on*_ and *k*_*off*_ were calculated as 21.7±13.8×10^8^ M^-1^×s^-1^ and 7.32±4.95×10^−6^ s^-1^, being highly consistent with the corresponding experimental values^32^ of 6.0×10^8^ M^-1^×s^-1^ and 8.0×10^−6^ s^-1^, respectively (**Table 2**).

## Discussion

We have developed a new “PPI-GaMD” computational approach to efficient enhanced sampling and accurate calculations of protein-protein binding thermodynamics and kinetics. PPI-GaMD works by selectively boosting the interaction potential energy between protein binding partners. Microsecond-timescale PPI-GaMD simulations have allowed us to capture repetitive protein dissociation and binding as demonstrated on the barstar-barnase model system, which thus enabled accurate free energy and kinetics calculations of protein binding.

PPI-GaMD simulations revealed that electrostatic interactions played a critical role in barstar binding to barnase, being consistent with previous experimental^3, 28b^ and computational studies.^17a, 19b, 21^ The important role of electrostatic interactions were identified in earlier cMD simulations^17a^ and mutation experiments.^3^ Multiple hundreds-of-nanosecond cMD simulations captured binding of barstar to barnase^17a^ and suggested that electrostatic interactions played a major role in the protein diffusion. Schreiberet et al.^3^ combined mutation and binding experiments to identify the importance of electrostatic interactions in the barstar binding affinity and kinetics. This study, 2 μs PPI-GaMD simulations have captured both barstar binding and unbinding, further supporting the important role of electrostatic interactions in forming the “intermediate” and “bound” states of the barstar-barnase system (**Figs. 2,4 & S2**). The charged residue pairs, including the BN:K27-BS:E80, BN:R59-BS:D39, BN:K27:BS:D39, BN:R83-BS:D39 and BN:R87-BS:D39, played a critical role in barstar binding and unbinding kinetics, thus effecting their binding thermodynamics, being consistent with previous mutation experiments^28b^. The k_on_ and k_off_ of wildtype barnase-barstar were changed from (3.7*10^8^ s^-1^M^-1^, 3.7*10^−6^ s^-1^) to (0.32*10^8^ s^-1^M^-1^, 3500*10^−6^ s^-1^), (0.58*10^8^ s^-1^M^-1^, 68000*10^−6^ s^-1^),(1.0*10^8^ s^-1^M^-1^, 53000*10^−6^ s^-1^), (1.6*10^8^ s^-1^M^-1^, 300000*10^−6^ s^-1^) in the double mutants of the BN:K27A/BS:E80A, BN:K27A/BS:D39A, BN:R83A/BS:D39A and BN:R87A/BS:D39A, respectively.^28b^ Therefore, the barstar binding free energy was changed from −19.0 kcal/mol in the wildtype to −13.5 kcal/mol, −10.8 kcal/mol, −12.6 kcal/mol, and -11.9 kcal/mol in the BN:K27A/BS:E80A, BN:K27A/BS:D39A, BN:R83A/BS:D39A and BN:R87A/BS:D39A double mutants, respectively.^28b^ Significant changes were also observed in the BN:R59A and BS:D39A single mutants,^28b^ suggesting the important role of these two residues.

Compared with existing methods, including the cMD,^17a^ Weighted Ensemble,^21^ MSM,^22^ Tempered Binding,^17b^ Replica Exchange MD,^15a^ Umbrella Sampling^19^ and Metadynamics,^16b, 20^ PPI-GaMD provides a more efficient and/or easier-to-use approach to simulation of protein-protein binding and dissociation. In particular, ∼440 μs cMD simulations were needed to capture binding of barstar to barnase in order to accurately predict the protein binding kinetics.^17b^ However, the slow protein unbinding process was still beyond the reach of cMD simulations. Even with the newly developed Tempered Binding technique, hundreds-of-microsecond simulations were needed to capture unbinding of barstar from barnase.^17b^ A total of ∼18 μs cMD simulations were able to predict accurate protein binding rate constants (*k*_*on*_) with the Weighted Ensemble approach.^21^ However, the Weighted Ensemble simulations were still not able to model the slow protein unbinding.^21^ MSM was able to predict intermediate structures, energetics and kinetics of barnase-barstar binding by Plattner et al.^22^ However, a total of ∼2 ms simulations were needed to build the MSM. For the Replica Exchange method,^15a, 43^ a large number of replica simulations were often needed to model PPIs. Twelve replicas distributed in the temperature range of 290–620 K were needed in PTMetaD-WTE, which combined replica exchange and Metadynamics, to investigate the binding mechanism of the insulin-dimer.^43^ The PTMetaD-WTE simulations also required predefined, carefully chosen CVs with expert knowledge of the studied system. The predefined CVs could potentially lead to certain constraints on the protein binding pathway and conformational space. Such simulations could also suffer from the “hidden energy barrier” problem and slow convergence if important CVs were missing.^23a, 23g^ Overall, the above methods appeared computationally expensive, requiring tens to hundreds-of-microsecond simulations to characterize protein binding thermodynamics and kinetics. In this context, PPI-GaMD that has allowed us to capture repetitive protein binding and unbinding through microsecond-timescale simulations provides a highly efficient approach to enhanced sampling of protein binding and dissociation.

PPI-GaMD shall be of wide applicability for PPIs other than barstar binding to barnase. In addition to protein binding free energy, it has enabled calculations of protein dissociation and binding rate constants. Remarkably, the barstar dissociation from barnase was accelerated by ∼11-13 orders of magnitude. It is worth noting that slight decrease in the protein association rate (∼2.7 times) was observed in the PPI-GaMD simulations (**Table 2**), which was similar to our recently developed LiGaMD^31^ and Pep-GaMD^32^ methods. Nevertheless, PPI-GaMD could be further combined with other enhanced sampling methods to facilitate the protein rebinding in future studies, including the Replica Exchange^44^ and Weighted Ensemble^45^ that have been successfully combined with GaMD. On the other hand, coarse-grained modeling,^46^ which significantly reduces the system degrees of freedom, could be applied to extend simulation timescales and investigate PPIs such as binding of the G proteins to GPCRs.^47^ These techniques could be incorporated to further improve the PPI-GaMD efficiency and advance studies in enhanced sampling of PPIs.

## Acknowledgements

We appreciate the help of Prof. David Case for accessing the AMBER git repository to develop our new simulation algorithms. This work used supercomputing resources with allocation award TG-MCB180049 through the Extreme Science and Engineering Discovery Environment (XSEDE), which is supported by National Science Foundation grant number ACI-1548562, and project M2874 through the National Energy Research Scientific Computing Center (NERSC), which is a U.S. Department of Energy Office of Science User Facility operated under Contract No. DE-AC02-05CH11231, and the Research Computing Cluster at the University of Kansas. This work was supported in part by the National Institutes of Health (R01GM132572), and the startup funding in the College of Liberal Arts and Sciences at the University of Kansas.

## Supporting Information

A detailed description energetic reweighting methods, PPI-GaMD in AMBER, **Table S1** and **Figures S1**-**S2** are provided in the Supporting Information. This information is available free of charge via the Internet at http://pubs.acs.org.

## TOC graphic

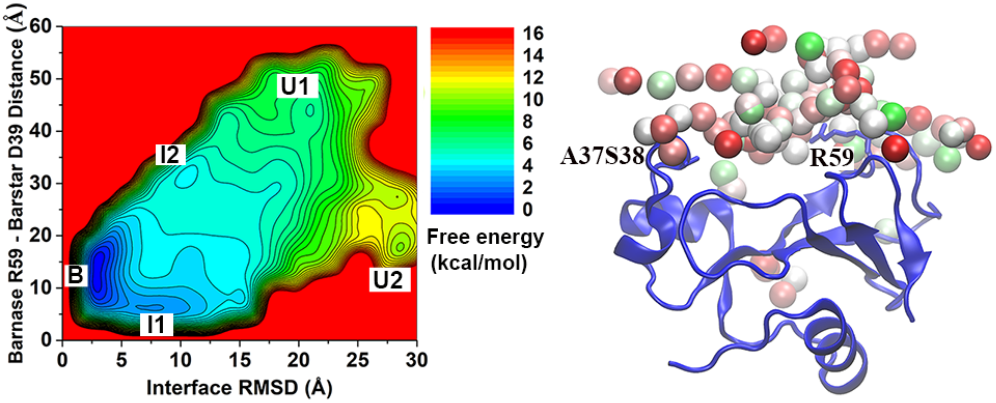

All-atom protein-protein interaction-Gaussian accelerated molecular dynamics (PPI-GaMD) simulations captured repeptitive binding and dissociaiton of barstar to the barnase model system, which enabled us to characterize the protein binding pathways, thermodynamcis and kinetics.

## References

1. Nooren, I. M.; Thornton, J. M., Diversity of protein-protein interactions. EMBO J. 2003, 22 (14), 3486–92.

2. (a)Scott, D. E.; Bayly, A. R.; Abell, C.; Skidmore, J., Small molecules, big targets: drug discovery faces the protein-protein interaction challenge. Nat. Rev. Drug Discov. 2016, 15 (8), 533–50; (b)Ferreira, L. G.; Oliva, G.; Andricopulo, A. D., Protein-protein interaction inhibitors: advances in anticancer drug design. Expert Opin. Drug Discov. 2016, 11 (10), 957–68.

3. Schreiber, G.; Fersht, A. R., Interaction of barnase with its polypeptide inhibitor barstar studied by protein engineering. Biochemistry 1993, 32 (19), 5145–50.

4. Miura, K., An Overview of Current Methods to Confirm Protein-Protein Interactions. Protein Pept. Lett. 2018, 25 (8), 728–733.

5. Sussman, J. L.; Lin, D.; Jiang, J.; Manning, N. O.; Prilusky, J.; Ritter, O.; Abola, E. E., Protein Data Bank (PDB): database of three-dimensional structural information of biological macromolecules. Acta Crystallogr., Sect. D: Biol. Crystallogr. 1998, 54 (Pt 6 Pt 1), 1078–84.

6. Vakser, I. A., Challenges in protein docking. Curr. Opin. Struct. Biol. 2020, 64, 160–165.

7. Karplus, M.; McCammon, J. A., Molecular dynamics simulations of biomolecules. Nat. Struct. Biol. 2002, 9 (9), 646–52.

8. Yan, Y.; Tao, H.; He, J.; Huang, S. Y., The HDOCK server for integrated protein-protein docking. Nat. Protoc. 2020, 15 (5), 1829–1852.

9. Tovchigrechko, A.; Vakser, I. A., GRAMM-X public web server for protein-protein docking. Nucleic Acids Res. 2006, 34 (Web Server issue), W310–4.

10. Park, T.; Woo, H.; Baek, M.; Yang, J.; Seok, C., Structure prediction of biological assemblies using GALAXY in CAPRI rounds 38-45. Proteins 2020, 88 (8), 1009–1017.

11. de Vries, S. J.; van Dijk, M.; Bonvin, A. M., The HADDOCK web server for data-driven biomolecular docking. Nat. Protoc. 2010, 5 (5), 883–97.

12. Li, X.; Moal, I. H.; Bates, P. A., Detection and refinement of encounter complexes for protein-protein docking: taking account of macromolecular crowding. Proteins 2010, 78 (15), 3189–96.

13. Kozakov, D.; Hall, D. R.; Xia, B.; Porter, K. A.; Padhorny, D.; Yueh, C.; Beglov, D.; Vajda, S., The ClusPro web server for protein-protein docking. Nat. Protoc. 2017, 12 (2), 255–278.

14. Huang, S. Y.; Zou, X., MDockPP: A hierarchical approach for protein-protein docking and its application to CAPRI rounds 15-19. Proteins 2010, 78 (15), 3096–103.

15. (a)Siebenmorgen, T.; Zacharias, M., Efficient Refinement and Free Energy Scoring of Predicted Protein-Protein Complexes Using Replica Exchange with Repulsive Scaling. J. Chem. Inf. Model. 2020, 60 (11), 5552–5562; (b)Siebenmorgen, T.; Engelhard, M.; Zacharias, M., Prediction of protein-protein complexes using replica exchange with repulsive scaling. J. Comput. Chem. 2020, 41 (15), 1436–1447.

16. (a)Zhang, L.; Borthakur, S.; Buck, M., Dissociation of a Dynamic Protein Complex Studied by All-Atom Molecular Simulations. Biophys. J. 2016, 110 (4), 877–86; (b)Banerjee, P.; Bagchi, B., Dynamical control by water at a molecular level in protein dimer association and dissociation. Proc. Natl. Acad. Sci. USA 2020, 117 (5), 2302–2308.

17. (a)Ahmad, M.; Gu, W.; Geyer, T.; Helms, V., Adhesive water networks facilitate binding of protein interfaces. Nat. Commun. 2011, 2, 261; (b)Pan, A. C.; Jacobson, D.; Yatsenko, K.; Sritharan, D.; Weinreich, T. M.; Shaw, D. E., Atomic-level characterization of protein-protein association. Proc. Natl. Acad. Sci. USA 2019, 116 (10), 4244–4249.

18. Kingsley, L. J.; Esquivel-Rodriguez, J.; Yang, Y.; Kihara, D.; Lill, M. A., Ranking protein-protein docking results using steered molecular dynamics and potential of mean force calculations. J. Comput. Chem. 2016, 37 (20), 1861–5.

19. (a)Gumbart, J. C.; Roux, B.; Chipot, C., Efficient determination of protein-protein standard binding free energies from first principles. J. Chem. Theory Comput. 2013, 9 (8), 3789–3798; (b)Joshi, D. C.; Lin, J. H., Delineating Protein-Protein Curvilinear Dissociation Pathways and Energetics with Naive Multiple-Walker Umbrella Sampling Simulations. J. Comput. Chem. 2019, 40 (17), 1652–1663.

20. Antoszewski, A.; Feng, C. J.; Vani, B. P.; Thiede, E. H.; Hong, L.; Weare, J.; Tokmakoff, A.; Dinner, A. R., Insulin Dissociates by Diverse Mechanisms of Coupled Unfolding and Unbinding. J. Phys. Chem. B 2020, 124 (27), 5571–5587.

21. Saglam, A. S.; Chong, L. T., Protein-protein binding pathways and calculations of rate constants using fully-continuous, explicit-solvent simulations. Chem. Sci. 2019, 10 (8), 2360–2372.

22. Plattner, N.; Doerr, S.; De Fabritiis, G.; Noe, F., Complete protein-protein association kinetics in atomic detail revealed by molecular dynamics simulations and Markov modelling. Nat. Chem. 2017, 9 (10), 1005–1011.

23. (a)Abrams, C.; Bussi, G., Enhanced Sampling in Molecular Dynamics Using Metadynamics, Replica-Exchange, and Temperature-Acceleration. Entropy 2014, 16 (1), 163–199; (b)Spiwok, V.; Sucur, Z.; Hosek, P., Enhanced sampling techniques in biomolecular simulations. Biotechnol. Adv. 2015, 33 (6 Pt 2), 1130–40; (c)Dellago, C.; Bolhuis, P. G., Transition Path Sampling and Other Advanced Simulation Techniques for Rare Events. Advanced Computer Simulation Approaches for Soft Matter Sciences Iii 2009, 221, 167–233; (d)Gao, Y. Q.; Yang, L. J.; Fan, Y. B.; Shao, Q., Thermodynamics and kinetics simulations of multi-time-scale processes for complex systems. Int. Rev. Phys. Chem. 2008, 27 (2), 201–227; (e)Liwo, A.; Czaplewski, C.; Oldziej, S.; Scheraga, H. A., Computational techniques for efficient conformational sampling of proteins. Curr. Opin. Struct. Biol. 2008, 18 (2), 134–9; (f)Christen, M.; Van Gunsteren, W. F., On searching in, sampling of, and dynamically moving through conformational space of biomolecular systems: A review. J. Comput. Chem. 2008, 29 (2), 157–166; (g)Miao, Y.; McCammon, J. A., Unconstrained Enhanced Sampling for Free Energy Calculations of Biomolecules: A Review. Mol. Simul. 2016, 42 (13), 1046–1055.

24. (a)Wang, J. N.; Arantes, P. R.; Bhattarai, A.; Hsu, R. V.; Pawnikar, S.; Huang, Y. M. M.; Palermo, G.; Miao, Y. L., Gaussian accelerated molecular dynamics: Principles and applications. Wiley Interdiscip. Rev.: Comput. Mol. Sci. 2021, 11 (5), e1521; (b)Miao, Y.; Feher, V. A.; McCammon, J. A., Gaussian Accelerated Molecular Dynamics: Unconstrained Enhanced Sampling and Free Energy Calculation. J. Chem. Theory Comput. 2015, 11 (8), 3584–3595.

25. (a)Pang, Y. T.; Miao, Y.; Wang, Y.; McCammon, J. A., Gaussian Accelerated Molecular Dynamics in NAMD. J. Chem. Theory Comput. 2017, 13 (1), 9–19; (b)Miao, Y.; McCammon, J. A., Gaussian Accelerated Molecular Dynamics: Theory, Implementation, and Applications. Annu. Rep. Comput. Chem. 2017, 13, 231–278; (c)Miao, Y., Acceleration of biomolecular kinetics in Gaussian accelerated molecular dynamics. J. Chem. Phys. 2018, 149 (7), 072308.

26. Miao, Y.; McCammon, J. A., Mechanism of the G-protein mimetic nanobody binding to a muscarinic G-protein-coupled receptor. Proc. Natl. Acad. Sci. USA 2018, 115 (12), 3036–3041.

27. Paul, F.; Wehmeyer, C.; Abualrous, E. T.; Wu, H.; Crabtree, M. D.; Schoneberg, J.; Clarke, J.; Freund, C.; Weikl, T. R.; Noe, F., Protein-peptide association kinetics beyond the seconds timescale from atomistic simulations. Nat. Commun. 2017, 8 (1), 1095.

28. (a)Frisch, C.; Schreiber, G.; Johnson, C. M.; Fersht, A. R., Thermodynamics of the interaction of barnase and barstar: changes in free energy versus changes in enthalpy on mutation. J. Mol. Biol. 1997, 267 (3), 696–706; (b)Schreiber, G.; Fersht, A. R., Energetics of protein-protein interactions: Analysis ofthe Barnase-Barstar interface by single mutations and double mutant cycles. J. Mol. Biol. 1995, 248 (2), 478–486.

29. (a)Vanommeslaeghe, K.; MacKerell, A. D., Jr., CHARMM additive and polarizable force fields for biophysics and computer-aided drug design. Biochim. Biophys. Acta, Gen. Subj. 2015, 1850 (5), 861–871; (b)Duan, Y.; Wu, C.; Chowdhury, S.; Lee, M. C.; Xiong, G.; Zhang, W.; Yang, R.; Cieplak, P.; Luo, R.; Lee, T.; Caldwell, J.; Wang, J.; Kollman, P., A point-charge force field for molecular mechanics simulations of proteins based on condensed-phase quantum mechanical calculations. J. Comput. Chem. 2003, 24 (16), 1999–2012.

30. Case, D. A. D.S. Cerutti; T.E. Cheatham, I. T.A. Darden, R. E. D., T.J. Giese, H. Gohlke, A.W. Goetz, D. Greene, N. Homeyer, S. Izadi, A. Kovalenko, T.S. Lee, S. LeGrand, P. Li, C. Lin, J. Liu, T. Luchko, R. Luo, D. Mermelstein, K.M. Merz, G. Monard, H. Nguyen, I. Omelyan, A. Onufriev, F. Pan, R. Qi, D.R. Roe, A. Roitberg, C. Sagui, C.L. Simmerling, W.M. Botello-Smith, J. Swails, R.C. Walker, J. Wang, R.M. Wolf, X. Wu, L. Xiao, D.M. York and P.A. Kollman, AMBER 2020, University of California, San Francisco. 2020.

31. Miao, Y.; Bhattarai, A.; Wang, J., Ligand Gaussian Accelerated Molecular Dynamics (LiGaMD): Characterization of Ligand Binding Thermodynamics and Kinetics. J. Chem. Theory Comput. 2020, 16 (9), 5526–5547.

32. Wang, J.; Miao, Y., Peptide Gaussian accelerated molecular dynamics (Pep-GaMD): Enhanced sampling and free energy and kinetics calculations of peptide binding. J. Chem. Phys. 2020, 153 (15), 154109.

33. (a)Doudou, S.; Burton, N. A.; Henchman, R. H., Standard Free Energy of Binding from a One-Dimensional Potential of Mean Force. J. Chem. Theory Comput. 2009, 5 (4), 909–18; (b)Buch, I.; Giorgino, T.; De Fabritiis, G., Complete reconstruction of an enzyme-inhibitor binding process by molecular dynamics simulations. Proc. Natl. Acad. Sci. USA 2011, 108 (25), 10184–9.

34. Miao, Y.; Sinko, W.; Pierce, L.; Bucher, D.; Walker, R. C.; McCammon, J. A., Improved Reweighting of Accelerated Molecular Dynamics Simulations for Free Energy Calculation. J. Chem. Theory Comput. 2014, 10 (7), 2677–2689.

35. (a)Doshi, U.; Hamelberg, D., Extracting Realistic Kinetics of Rare Activated Processes from Accelerated Molecular Dynamics Using Kramers’ Theory. J. Chem. Theory Comput. 2011, 7 (3), 575–81; (b)Frank, A. T.; Andricioaei, I., Reaction Coordinate-Free Approach to Recovering Kinetics from Potential-Scaled Simulations: Application of Kramers’ Rate Theory. J. Phys. Chem. B 2016, 120 (33), 8600–5.

36. Hamelberg, D.; Shen, T.; Andrew McCammon, J., Relating kinetic rates and local energetic roughness by accelerated molecular-dynamics simulations. J. Chem. Phys. 2005, 122 (24), 241103.

37. Buckle, A. M.; Schreiber, G.; Fersht, A. R., Protein-protein recognition: crystal structural analysis of a barnase-barstar complex at 2.0-A resolution. Biochemistry 1994, 33 (30), 8878–89.

38. Maier, J. A.; Martinez, C.; Kasavajhala, K.; Wickstrom, L.; Hauser, K. E.; Simmerling, C., ff14SB: Improving the Accuracy of Protein Side Chain and Backbone Parameters from ff99SB. J. Chem. Theory Comput. 2015, 11 (8), 3696–713.

39. Jorgensen, W. L.; Chandrasekhar, J.; Madura, J. D.; Impey, R. W.; Klein, M. L., Comparison of Simple Potential Functions for Simulating Liquid Water. J. Chem. Phys. 1983, 79 (2), 926–935.

40. Humphrey, W.; Dalke, A.; Schulten, K., VMD: visual molecular dynamics. J. Mol. Graph. 1996, 14 (1), 33-8, 27-8.

41. Roe, D. R.; Cheatham, T. E., 3rd, PTRAJ and CPPTRAJ: Software for Processing and Analysis of Molecular Dynamics Trajectory Data. J. Chem. Theory Comput. 2013, 9 (7), 3084–95.

42. Ribeiro, J. M. L.; Tsai, S. T.; Pramanik, D.; Wang, Y.; Tiwary, P., Kinetics of Ligand-Protein Dissociation from All-Atom Simulations: Are We There Yet? Biochemistry 2019, 58 (3), 156–165.

43. Banerjee, P.; Mondal, S.; Bagchi, B., Insulin dimer dissociation in aqueous solution: A computational study of free energy landscape and evolving microscopic structure along the reaction pathway. J. Chem. Phys. 2018, 149 (11), 114902.

44. Oshima, H.; Re, S.; Sugita, Y., Replica-Exchange Umbrella Sampling Combined with Gaussian Accelerated Molecular Dynamics for Free-Energy Calculation of Biomolecules. J. Chem. Theory Comput. 2019, 15 (10), 5199–5208.

45. Ahn, S.-H.; Ojha, A.; Amaro, R.; McCammon, J., Gaussian accelerated molecular dynamics with the weighted ensemble method: a hybrid method improves thermodynamics and kinetics sampling. ChemRxiv 2021.

46. (a)Lamprakis, C.; Andreadelis, I.; Manchester, J.; Velez-Vega, C.; Duca, J. S.; Cournia, Z., Evaluating the Efficiency of the Martini Force Field to Study Protein Dimerization in Aqueous and Membrane Environments. J. Chem. Theory Comput. 2021, 17 (5), 3088–3102; (b)Dhusia, K.; Su, Z. Q.; Wu, Y. H., Using Coarse-Grained Simulations to Characterize the Mechanisms of Protein-Protein Association. Biomolecules 2020, 10 (7), 1056.

47. Mondal, D.; Kolev, V.; Warshel, A., Exploring the activation pathway and Gi-coupling specificity of the mu-opioid receptor. Proc. Natl. Acad. Sci. USA 2020, 117 (42), 26218–26225.

