## Supplementary material for "Protein-protein interaction-Gaussian accelerated molecular dynamics (PPI-GaMD): Characterization of protein binding thermodynamics and kinetics": SI

### Methods

#### Energetic reweighting

Energetic reweighting of PPI-GaMD simulations is similar to that of previous GaMD[1] for calculating the potential of mean force (PMF). The probability distribution along a reaction coordinate is written as $p^*(A)$ . Given the boost potential $\Delta V(r)$ of each frame, $p^*(A)$ can be reweighted to recover the canonical ensemble distribution, $p(A)$, as:

$$p(A_j) = p^*(A_j)\frac{\langle e^{\beta \Delta V(r)}\rangle_j}{\sum_{i=1}^{M}\langle p^*(A_i) e^{\beta \Delta V(r)}\rangle_i}, \ j = 1, \ldots, M, \tag{S1}$$

where $M$ is the number of bins, $\beta = k_B T$ and $\langle e^{\beta \Delta V(r)}\rangle_j$ is the ensemble-averaged Boltzmann factor of $\Delta V(r)$ for simulation frames found in the $j^{\text{th}}$ bin. The ensemble-averaged reweighting factor can be approximated using cumulant expansion:

$$\langle e^{\beta \Delta V(r)}\rangle = exp\left\{\sum_{k=1}^{\infty}\frac{\beta^k}{k!}C_k\right\}, \tag{S2}$$

where the first two cumulants are given by:

$$\begin{aligned} C_1 &= \langle \Delta V\rangle, \\ C_2 &= \langle \Delta V^2\rangle - \langle \Delta V\rangle^2 = \sigma_v^2. \end{aligned} \tag{S3}$$

The boost potential obtained usually exhibits near-Gaussian distribution[1]. Cumulant expansion to the second order thus provides a good approximation for computing the reweighting factor.[2-3] The reweighted free energy $F(A) = -k_B T \ln p(A)$ is calculated as:

$$F(A) = F^*(A) - \sum_{k=1}^{2}\frac{\beta^k}{k!}C_k + F_c, \tag{S4}$$

where $F^*(A) = -k_B T \ln p^*(A)$ is the modified free energy obtained from PPI-GaMD simulation and $F_c$ is a constant.

#### Implementation of protein-protein interaction Gaussian accelerated molecular dynamics (PPI-GaMD)

PPI-GaMD is currently implemented in the GPU version of AMBER 20,[4] but should be transferable to other molecular dynamics programs as well. PPI-GaMD provides enhanced sampling of protein-protein interactions (PPIs). Following is a list of the input parameters for a PPI-GaMD simulation:

***igamd*** Flag to apply boost potential
= **0** (default) no boost is applied
= **1** boost on the total potential energy only (GaMD_Tot)
= **2** boost on the dihedral energy only (GaMD_Dih)
= **3** dual boost on both dihedral and total potential energy (GaMD_Dual)
= **4** boost on the non-bonded potential energy only (GaMD_NB)
= **5** dual boost on both dihedral and non-bonded potential energy (GaMD_NB_Dual)
= **10** boost on non-bonded potential energy of selected region (defined by timask1 and scmask1) as for a ligand (LiGaMD)

= **11** dual boost on both non-bonded potential energy of the bound ligand and the remaining potential energy of the entire system (LiGaMD_Dual)
= **14** boost on the total potential energy of selected region (defined by timask1 and scmask1) as for a peptide (Pep-GaMD)
= **15** dual boost on both the peptide essential interaction potential energy and the remaining potential energy of the entire system (Pep-GaMD_Dual)
= **16** boost on the interaction between protein partners (The first protein is defined by timask1 and scmask1 and the second one defined by bgpro2atm (first atom number of the protein) and edpro2atm (the end atom number of the protein)) for protein-protein interaction GaMD (PPI-GaMD)
= **17** dual boost on both the protein protein interactions and the remaining potential energy of the entire system (PPI-GaMD_Dual)

***iE*** Flag to set the threshold energy $E$ for applying all boost potentials
= **1** (default) set the threshold energy to the lower bound $E = V_{\max}$
= **2** set the threshold energy to the upper bound $E = V_{\min} + (V_{\max} - V_{\min})/k_0$

***iEP*** Flag to overwrite *iE* and set the threshold energy $E$ for applying the first boost potential in dual-boost schemes
= **1** (default) set the threshold energy to the lower bound $E = V_{\max}$
= **2** set the threshold energy to the upper bound $E = V_{\min} + (V_{\max} - V_{\min})/k_0$

***iED*** Flag to overwrite *iE* and set the threshold energy $E$ for applying the second boost potential in dual-boost schemes
= **1** (default) set the threshold energy to the lower bound $E = V_{\max}$
= **2** set the threshold energy to the upper bound $E = V_{\min} + (V_{\max} - V_{\min})/k_0$

***ntcmdprep*** The number of preparation conventional molecular dynamics steps. This is used for system equilibration and the potential energies are not collected for calculating their statistics. The default is 200,000 for a simulation with 2 *fs* timestep.

***ntcmd*** The number of initial conventional molecular dynamics simulation steps used to calculate the maximum, minimum, average and standard deviation of the system potential energies (i.e., $V_{\max}$, $V_{\min}$, $V_{\text{avg}}$, $\sigma_V$). The default is 1,000,000 for a simulation with 2 *fs* timestep.

***ntebprep*** The number of preparation biasing molecular dynamics simulation steps. This is used for system equilibration after adding the boost potential and the potential statistics (i.e., $V_{\max}$, $V_{\min}$, $V_{\text{avg}}$, $\sigma_V$) are not updated during these steps. The default is 200,000 for a simulation with 2 fs timestep.

***nteb*** The number of biasing molecular dynamics simulation steps. Potential statistics ($V_{\max}$, $V_{\min}$, $V_{\text{avg}}$, $\sigma_V$) are updated between the ***ntebprep*** and ***nteb*** steps and used to calculate the GaMD acceleration parameters, particularly $E$ and $k_0$. The default is 1,000,000 for a simulation with 2 *fs* timestep. A greater value may be needed to ensure that the potential statistics and GaMD acceleration parameters level off before running production simulation between the ***nteb*** and ***nstlim*** (total simulation length) steps. Moreover, ***nteb*** can be set to ***nstlim***, by which the potential statistics and GaMD acceleration parameters are updated adaptively throughout the simulation. This in some cases provides more appropriate acceleration.

***ntave*** The number of simulation steps used to calculate the average and standard deviation of potential energies. This variable has already been used in AMBER. The default is set to 50,000 for GaMD simulations. It is recommended to be updated as about 4 times of the total number of atoms in the system. Note that ***ntcmd*** and ***nteb*** need to be multiples of ***ntave***.

***irest_gamd*** Flag to restart GaMD simulation

**= 0** (default) new simulation. A file "gamd-restart.dat" that stores the maximum, minimum, average and standard deviation of the potential energies needed to calculate the boost potentials (depending on the ***igamd*** flag) will be saved automatically after GaMD equilibration stage.

**= 1** restart simulation (***ntcmd*** and ***nteb*** are set to 0 in this case). The "gamd-restart.dat" file will be read for restart.

***sigma0P*** The upper limit of the standard deviation of the first potential boost that allows for accurate reweighting. The default is 6.0 (unit: kcal/mol).

***sigma0D*** The upper limit of the standard deviation of the second potential boost that allows for accurate reweighting in dual-boost simulations (e.g., ***igamd*** = 2, 3, 5, 11, 15 and 17). The default is 6.0 (unit: kcal/mol).

***timask1*** Specifies atoms of the bound ligand in ambmask format. This variable has already been used in AMBER. The default is an empty string.

***scmask1*** Specifies atoms of the bound ligand that will be described using soft core in ambmask format. This variable has already been used in AMBER. The default is an empty string.

***bgpro2atm*** Start atomic number of the second protein.

***edpro2atm*** The final atomic number of the second protein.

Example input parameters used in PPI-GaMD simulations include the following:

```
icfe = 1, ifsc = 1, gti_cpu_output = 0,gti_add_sc = 1,
timask1 = ':1-110', scmask1 = ':1-110',
timask2 = '', scmask2 = '',
bgpro2atm=1, edpro2atm=1453,
igamd = 17, iEP = 2, iED = 1, irest_gamd = 0,
ntcmd = 1000000, nteb = 1000000, ntave = 50000,
ntcmdprep = 200000, ntebprep = 200000,
sigma0P = 6.0, sigma0D = 6.0,
```

The PPI-GaMD algorithm is summarized as the following:

```
PPI-GaMD {
  If (irest_gamd == 0) then
    For i = 1, …, ntcmd // run initial conventional molecular dynamics
      If (i >= ntcmdprep) Update Vmax and Vmin of interaction potential energy
      If (i >= ntcmdprep && i%ntave ==0) Update Vavg and sigmaV of interaction potential energy
    End
   Save Vmax,Vmin,Vavg,sigmaV of interaction potential energy to "gamd_restart.dat" file
   Calc_E_k0(iE,sigma0,Vmax,Vmin,Vavg,sigmaV)
```

```
    For i = ntcmd+1, …, ntcmd+nteb // Run biasing molecular dynamics simulation steps
        deltaV = 0.5*k0*(E-V)**2/(Vmax-Vmin)
        V = V + deltaV
        If (i >= ntcmd+ntebprep) Update Vmax and Vmin of interaction potential energy
        If (i >= ntcmd+ntebprep && i%ntave ==0) Update Vavg and sigmaV of interaction potential energy
        Calc_E_k0(iE,sigma0,Vmax,Vmin,Vavg,sigmaV)
    End
    Save Vmax,Vmin,Vavg and sigmaV of of interaction potential energy to "gamd_restart.dat" file
  else if (irest_gamd == 1) then
    Read Vmax,Vmin,Vavg and sigmaV of interaction potential energy from "gamd_restart.dat" file
  End if


  For i = ntcmd+nteb+1, …, nstlim // run production simulation
    deltaV = 0.5*k0*(E-V)**2/(Vmax-Vmin)
    V = V + deltaV
  End
}

Subroutine Calc_E_k0(iE,sigma0,Vmax,Vmin,Vavg,sigmaV) {
if iE = 1 :
        E = Vmax
        k0' = (sigma0/sigmaV) * (Vmax-Vmin)/(Vmax-Vavg)
        k0 = min(1.0, k0')
else if iE = 2 :
        k0" = (1-sigma0/sigmaV) * (Vmax-Vmin)/(Vavg-Vmin)
        if 0 < k0" <= 1 :
                k0 = k0"
                E = Vmin + (Vmax-Vmin)/k0
        else
                E = Vmax
                k0' = (sigma0/sigmaV) * (Vmax-Vmin)/(Vmax-Vavg)
                k0 = min(1.0, k0')
        end
end
}
```

**Table S1.** Energy barriers of barnase-barstar dissociation ("off") and binding ("on") calculated from the reweighed (***ΔF***) and modified (no reweighting, ***ΔF****) free energy profiles, curvatures of the reweighed (*w*) and modified (*w*∗) free energy profiles near the guest Bound ("B"), Barrier ("Br") and Unbound ("U") states, and the ratio of apparent diffusion coefficients calculated from the PPI-GaMD simulations without reweighting (modified, *D*∗) and with reweighting (*D*). $k_{on}$ and $k_{off}$ are the kinetic dissociation and binding rate constants, respectively, from PPI-GaMD simulations with reweighting using Kramers' rate theory.

| Simulation | Δ***F*** **(kcal/mol)** | | Δ***F**** **(kcal/mol)** | | ***w*** | | ***w**** | | | | ***D*/D*** | | $k_{off}$ ($10^{-6} S^{-1}$) | $k_{on}$ ($10^{8} M^{-1}$) |
|---|---|---|---|---|---|---|---|---|---|---|---|---|---|---|
| | **Off** | **On** | **Off** | **On** | **B** | **Br** | **U** | **B** | **Br** | **U** | **Off** | **On** | | |
| Sim1 | 6.91 | 1.06 | 1.41 | 0.23 | 3.25 | 0.28 | 0.66 | 0.34 | 1.36 | 1.53 | 0.094 | 0.52 | 9.33 | 0.33 |
| Sim2 | 7.70 | 2.01 | 1.89 | 0.10 | 0.72 | 0.71 | 0.73 | 0.36 | 0.14 | 0.1 | 0.55 | 0.68 | 6.04 | 7.08 |
| Sim3 | 7.60 | 4.8 | 1.91 | 0.76 | 0.40 | 2.28 | 19.5 | 0.18 | 0.14 | 0.28 | 0.56 | 0.25 | 19.31 | 23.93 |
| Sim4 | 5.85 | 0.95 | 0.18 | 0.16 | 0.33 | 0.25 | 0.56 | 0.21 | 0.047 | 0.45 | 0.49 | 0.26 | 8.16 | 20.61 |
| Sim5 | 9.40 | 1.52 | 1.67 | 0.54 | 6.98 | 0.31 | 0.48 | 0.51 | 0.13 | 0.109 | 1.45 | 0.197 | 0.095 | 46.26 |
| Sim6 | 7.72 | 1.16 | 1.68 | 0.21 | 1.29 | 0.24 | 0.13 | 6.55 | 0.010 | 0.36 | 0.69 | 0.22 | 0.97 | 32.17 |

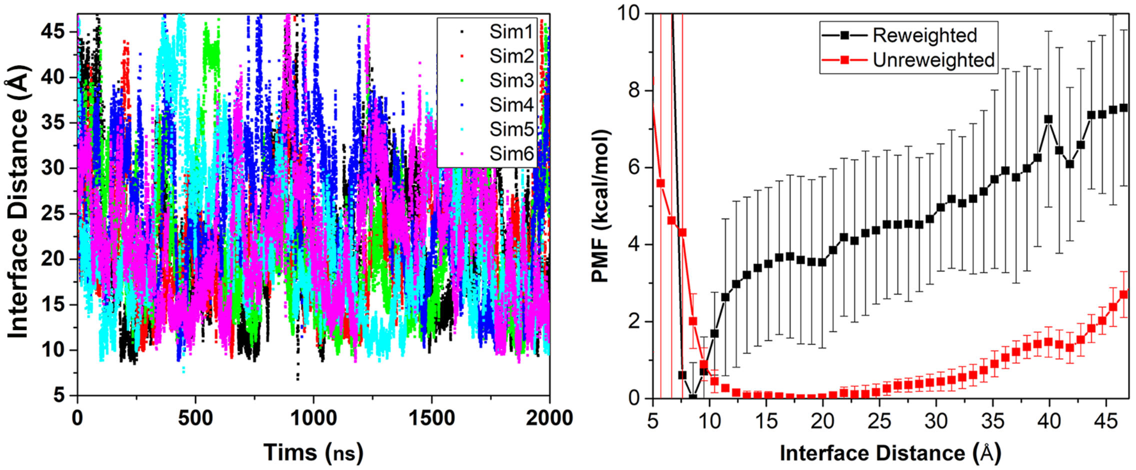


**Figure S1.** (**A**) Time courses of protein-protein interface distance calculated from six independent 2 μs PPI-GaMD simulations. (**B**) Original (reweighted) and modified (no reweighting) PMF profiles of the protein interface distance averaged over six PPI-GaMD simulations. Error bars are standard deviations of the free energy values calculated from six PPI-GaMD simulations.

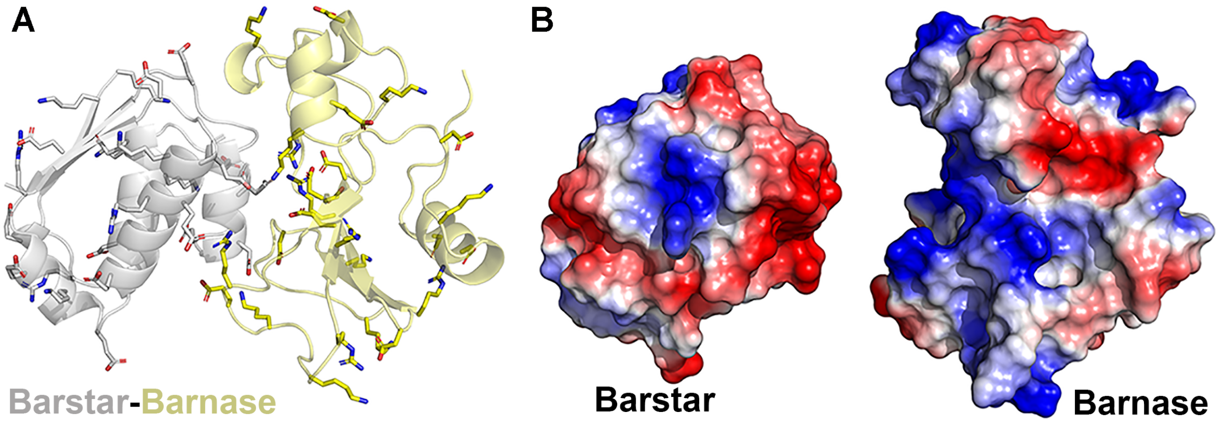


**Figure S2.** (**A**) Charged residues in barnase and barstar as highlighted in sticks. The nitrogen atoms in positively charged residues are colored blue. The oxygen atoms in negatively charged residues are colored red. (**B**) Electrostatic potential of the barstar and barnase. The interfacial distance in the crystal structure of the Barnase-Barstar complex (chain B and chain F in the 1BRS.pdb) was increased by 10 Å.

**References**

(1) Miao, Y.; McCammon, J. A., Gaussian Accelerated Molecular Dynamics: Theory, Implementation and Applications. *Annu. Rep. Comp. Chem.* **2017,** *13*, 231-278.
(2) Miao, Y.; Sinko, W.; Pierce, L.; Bucher, D.; McCammon, J. A., Improved reweighting of accelerated molecular dynamics simulations for free energy calculation. *J. Chem. Theory Comput.* **2014,** *10* (7), 2677–2689.
(3) Miao, Y.; Feher, V. A.; McCammon, J. A., Gaussian Accelerated Molecular Dynamics: Unconstrained Enhanced Sampling and Free Energy Calculation. *J. Chem. Theory Comput.* **2015,** *11* (8), 3584-3595.
(4) Case, D. A.; Belfon, K.; Ben-Shalom, I. Y.; Brozell, S. R.; Cerutti, D. S.; T.E. Cheatham, I.; Cruzeiro, V. W. D.; Darden, T. A.; Duke, R. E.; Giambasu, G., et al., AMBER 20, University of California, San Francisco. **2020**.
